# PERSISTENT RELIEF OF MOTOR SYMPTOMS IN A PARKINSONIAN MOUSE MODEL AFTER INDUCTION OF LTD AT CORTICAL INPUTS TO INDIRECT PATHWAY STRIATAL NEURONS

**DOI:** 10.1101/2022.01.21.477283

**Authors:** Chandrika Abburi, Jason Thome, Braeden Rodriguez, Xiaoxi Zhaung, Daniel S McGehee

## Abstract

In Parkinson’s disease (PD) patients, dopamine replacement therapy requires days to reach maximal effects, and the return of symptoms without treatment is similarly delayed. We previously postulated that these phenomena are mediated by plasticity of coritcostriatal synapses. As dopamine depletion is expected to promote aberrant potentiation of the cortical inputs onto indirect pathway neurons, we reasoned that induction of LTD here could reduce motor deficits in a PD model. Optogenetic cortical stimulation combined with a D2 receptor agonist, quinpirole, induces robust optical LTD (oLTD) in brain slices from 6-OHDA lesioned mice. When lesioned mice were subjected to corticostriatal oLTD treatment over 5 days, motor performance was improved for >3 weeks. Consistent with LTD induction, oLTD-treated mice had reduced VGLUT1 expression in striatum and greater excitability of D2 neurons. These findings suggest that reversing aberrant corticostriatal synaptic plasticity in the indirect pathway may lead to persistent relief of PD motor symptoms.

## Introduction

Dopamine (DA) in dorsal striatum is critical for motor learning and performance (Sommer et al., 2014). In the classic model, striatal DA release activates D1 receptors to increase excitability of direct pathway medium spiny neurons (D1-MSNs), while D2 receptor activation decreases activity of indirect pathway MSNs (D2-MSNs). Both these changes in excitability promote movement (Lahiri and Bevan, 2020; Nicola et al., 2000; Shen et al., 2008), however, decreased DA receptor activity following the loss of nigrostriatal DA neurons in Parkinson’s disease (PD) leads to the commonly observed motor deficits in those patients.

In addition to changes in excitability, existing evidence indicates that striatal DA also contributes to motor learning through its impact on cAMP levels and corticostriatal plasticity. DA promotes long-term plasticity of striatal glutamate synapses, specifically long-term potentiation (LTP) of inputs to D1-MSNs, and long-term depression (LTD) of the inputs to D2-MSNs (Augustin et al., 2014; Calabresi et al., 2007a, 2007b; Centonze et al., 2001; Gerfen and Surmeier, 2011; Shen et al., 2008). In one study, loss of LTD in the indirect pathway corticostriatial synapses contributes to motor deficits, whereas pharmacological interventions that rescue LTD also rescues motor performance (Kreitzer and Malenka, 2007).

Our previous findings demonstrated that loss of DA signaling through blockade of D2 receptors causes aberrant inhibitory motor learning and gradual deterioration of motor performance. Importantly, such aberrant inhibitory motor learning is experience-dependent and task-specific. Moreover, these motor deficits develop gradually, and they persist, even after restoration of DA neurotransmission. Those data suggest a key motor learning mechanism involving plasticity of corticostriatial synapses in the indirect pathway (Beeler et al., 2012).

When DA is depleted in PD patients or animal models, DA replacement therapy takes a few days to reach maximal therapeutic effects, while cessation of treatment results in persistent relief of symptoms that lasts for days or even weeks. This important phenomenon in DA replacement therapy is known as the ‘long-duration response’ (LDR) (Hauser et al., 2009; Nagao and Patel, 2019). In these patients, the slow onset of therapeutic effects and the persistent motor improvement suggest that DA replacement promotes striatal synaptic plasticity, particularly at the principal excitatory inputs from motor cortex.

Our model suggests that DA deficits promote aberrant LTP at the cortical inputs to D2-MSNs. As such, we tested whether reversing that plasticity using an optogenetic approach could rescue motor performance in a persistent manner, similar to the LDR. Our data suggest that high frequency motor cortical stimulation in the presence of a D2 agonist improves motor performance for over 4 weeks in a mouse model of PD.

## Results

### Combined optogenetic stimulation of motor cortical inputs to dorsolateral striatum with a D2 receptor agonist improves motor performance in PD mice

Loss of DA in dorsolateral striatum (DLS) alters synaptic plasticity in D2-MSNs, (Calabresi et al., 1997; Ryan et al., 2018; Trusel et al., 2015), which has been hypothesized to underlie aberrant motor learning and performance (Beeler et al., 2012). Here we attempt to reverse the aberrant plasticity of corticostriatal synapses on D2-MSNs in DA depleted mice by combining optogenetic high-frequency stimulation (oHFS) of motor cortical afferents with systemic injection of a D2 receptor agonist, quinpirole (oHFS/Quin treatment). To optogenetically excite the layer V of primary motor cortex (M1), we used adult Rbp-4 Cre^+/-^ and Drd2-EGFP^+/-^ double transgenic mice with stereotaxic injections of adeno-associated virus serotype 2 (AAV2) carrying a Cre recombinase-dependent channelrhodopsin (ChR2) with mCherry reporter (AAV2-EF1a-DIO-hChR2(H134R)-mCherry) (Fig 1a,b). Optical fibers were implanted near the dorsolateral border of the striatum to selectively stimulate M1 projections to DLS (Fig 1a).

**Figure 1.**
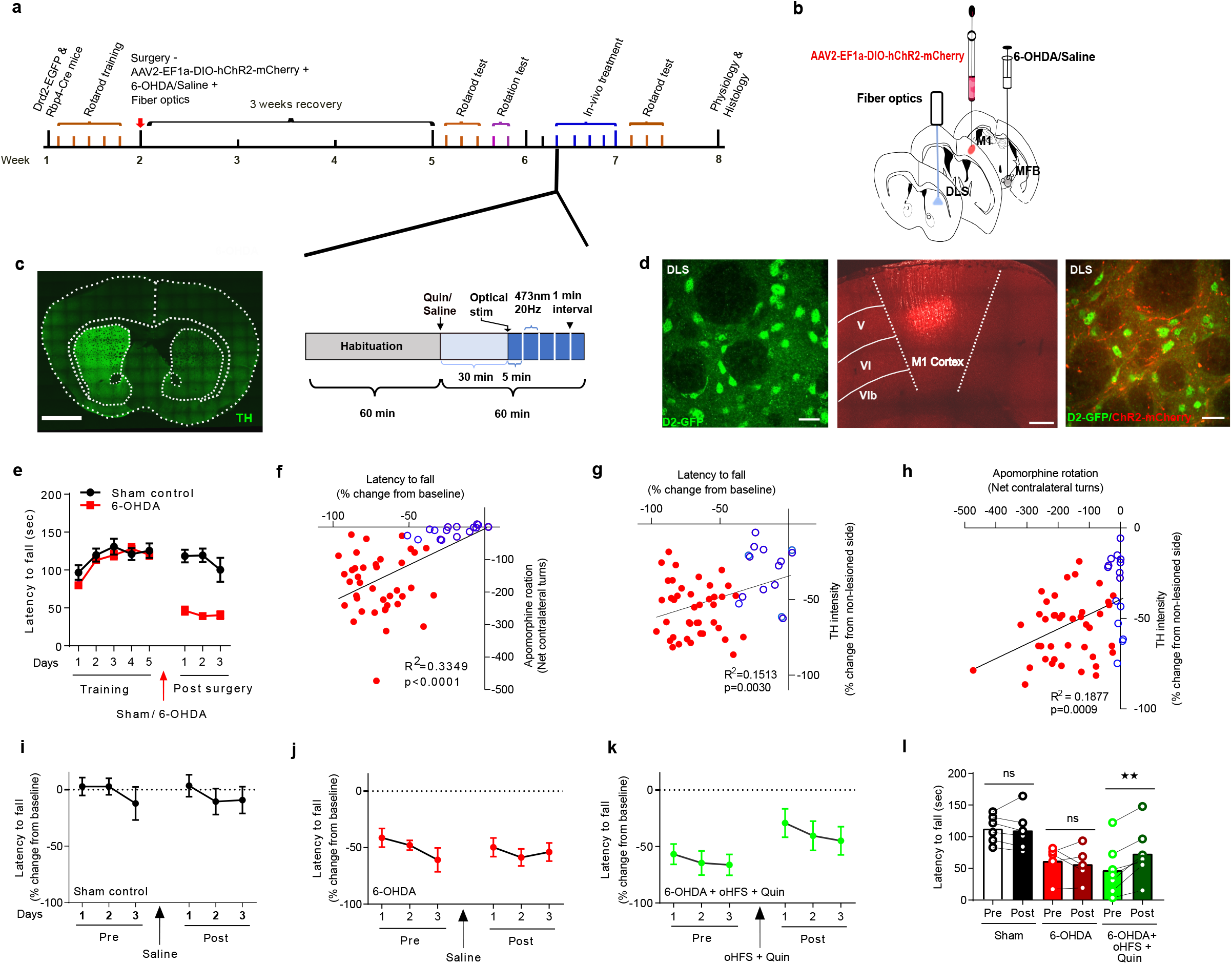
Motor improvement with D2 receptor agonist and oHFS treatment in 6-OHDA lesioned mice. (a) Experimental timeline for the behavioral assessment of 6-OHDA lesioned mice and *in-vivo* treatment strategy. (b) Schematic of 6-OHDA injection within MFB, AAV2-EF1a-DIO-ChR2-mCherry injection in M1 cortex of Rbp4-cre mice, and fiberoptic implantation location in DLS. (c) Coronal brain section with tyrosine hydroxylase (TH) immunostaining, confirming the loss of dopamine on lesioned side (right). (d) Representative coronal section from Drd2-EGFP mice showing EGFP fluorescence in D2-MSNs in DLS (left), layer V pyramidal neurons with ChR2 expression and mCherry fluorescence (middle), and cortical terminals with ChR2/mCherry fluorescence in proximity with D2-EGFP-expressing neurons DLS (right). (e) Rotarod performance in during 5 days of training and 3 weeks after 6-OHDA or sham surgery. Latency to fall from the accelerating rotarod is shown for sham controls (n=6), and 6-OHDA lesioned mice (n=36). (f) Linear regression correlation analysis between rotarod performance and apomorphine-induced rotation behavior. (g-h) Linear regression correlation analysis between rotarod or apomorphine-induced rotation behavior and TH fluorescence intensity. (i-k) Percent change in rotarod performance relative to pre-lesioned baseline before and after in vivo treatment. Sham (n=6), 6-OHDA with saline treatment (n=8) and 6-OHDA with quinpirole and optical stimulation (n=6). (l) Summary data of latency to fall from rotarod, showing individual changes in performance with in vivo treatments. Bars represent mean values for each condition. Scale bars: 1 mm in (c), 20 μm left and right and 500 μm middle in (d). Statistics: **p ≤ 0.005, paired t – test in (l).

To generate Parkinsonian mice, we injected 6-hydroxydopamine (6-OHDA) into the medial forebrain bundle unilaterally (Fig 1a, b), which resulted in depletion of DA in ipsilateral striatum, as shown by loss of tyrosine hydroxlase (TH) expression (Fig1c). In Drd2-EGFP mice, EGFP is expressed in D2 receptor-expressing neurons, allowing visualization in ex-vivo slice physiology (Fig 1d). Prior to stereotaxic surgeries, mice were trained on rotarod for 5 days to establish stable baseline performance (Fig 1e). Unilateral 6-OHDA injections into the median forebrain bundle or sham injections were performed, and after three weeks of recovery, the mice were tested for their motor performance on rotarod. To maximize the potential therapeutic effects of our treatment, mice with the strongest motor deficits were utilized for subsequent optogenetic stimulation. Motor deficits were assessed using rotarod and rotation behavioral tests to identify experimental subjects. Rotarod assessment demonstrated that 6-OHDA treatment decreased the latency to fall compared to baseline performance (Fig 1e). The apomorphine-induced rotation test involved challenging mice with apomorphine injections (0.5 mg/kg) to induce turning behavior contralateral to the lesioned brain hemisphere. Linear regression analysis revealed a strong positive correlation between these two behaviors (Fig 1f). Experimental subjects were identified as those displaying a 6-OHDA-induced change in rotarod performance greater that 30% relative to baseline, and >20 net turns contralateral to the lesioned hemisphere in a 30 min period after apomophine injection. A third level of assessment of the lesion involved post hoc TH immunostaining, where animals were excluded from the analysis if TH staining intensity was reduced by less than 20% of the non-lesioned side. Consistent with previous reports (O’Brien and Austin, 2019; Su et al., 2018) both these behavioral measures correlate with the 6-OHDA-induced reduction in TH intensity in DLS (Fig 1g, h).

Ex-vivo slice electrophysiology studies have shown that, high frequency stimulation (HFS) induces LTD at corticostriatal synapses onto D2-MSNs in unlesioned mice, however this form of plasticity is lost with DA depletion (Augustin et al., 2018; Skiteva et al., 2018; Trusel et al., 2015). Striatal HFS in the presence of D2 receptor activation rescues the plasticity at these synapses (Kitada et al., 2007). Based on these observations, we tested whether selective stimulation of M1 projections to DLS combined with D2 receptor activation would reverse motor deficits in PD mice. Thus, we examined the behavioral impact of optogenetic stimulation of M1 cortical afferents to DLS following systemic administration of the D2 receptor agonist, quinpirole. For these studies, lesioned mice that met the motor behavior criteria described above were subjected to oHFS/Quin stimulation sessions on 5 consecutive days (Fig 1a). Each session consisted of 60 min habituation followed by quinpirole administration (5 mg/kg, i.p.), and 30 min later, 5 trains of 20 Hz optical stimulation, 30 min each. The next day, rotarod testing resumed for 3 consecutive days. Control groups, either Sham 6-OHDA surgery (Fig 1i), or lesioned mice that received saline in place of oHFS/Quin treatment (Fig 1j) did not show altered motor performance compared to baseline. However, oHFS/Quin treatment significantly improved motor performance on rotarod in 6-OHDA lesioned mice (Fig 1k, l). Note that 7 of 7 subjects showed improved motor performance following oHFS/Quin treatment. These results suggest that optogenetic excitation of corticostriatal inputs to DLS in the presence of a D2 agonist improves motor performance in a mouse model of PD.

### Long lasting motor improvement in PD mice with combined optogenetic stimulation and quinpirole treatment

In PD patients, DA replacement therapies induce persistent improvement in motor function, even after cessation of treatment, which is known as the long duration response (Hauser et al., 2009; Nagao and Patel, 2019) Our group and others have suggested that this persistent effect is mediated by DA modulation of corticostriatal synaptic strength (Anderson et al., 2014; Beeler et al., 2012). Therefore, we investigated the duration of motor improvement following oHFS/Quin treatment. An independent cohort of mice was treated as illustrated in Fig 2a, and similar to the results in Fig 1, we observed improved motor performance in the days immediately after treatment (Fig 2b, 2e). We then retested their rotarod performance during the 3^rd^ and 4^th^ weeks after stimulation (Fig2b), and observed remarkably consistent performance into the 4^th^ week (Fig 2b, 2e). We next examined if subsequent oHFS/Quin treatment would further improve motor performance in these mice, and administered this during the 5^th^ week after the initial treatment. This additional treatment supported improved motor performance during the 6^th^ week, but there was no indication of additive effects relative to the first treatment (Fig 2b, 2e).

**Figure 2.**
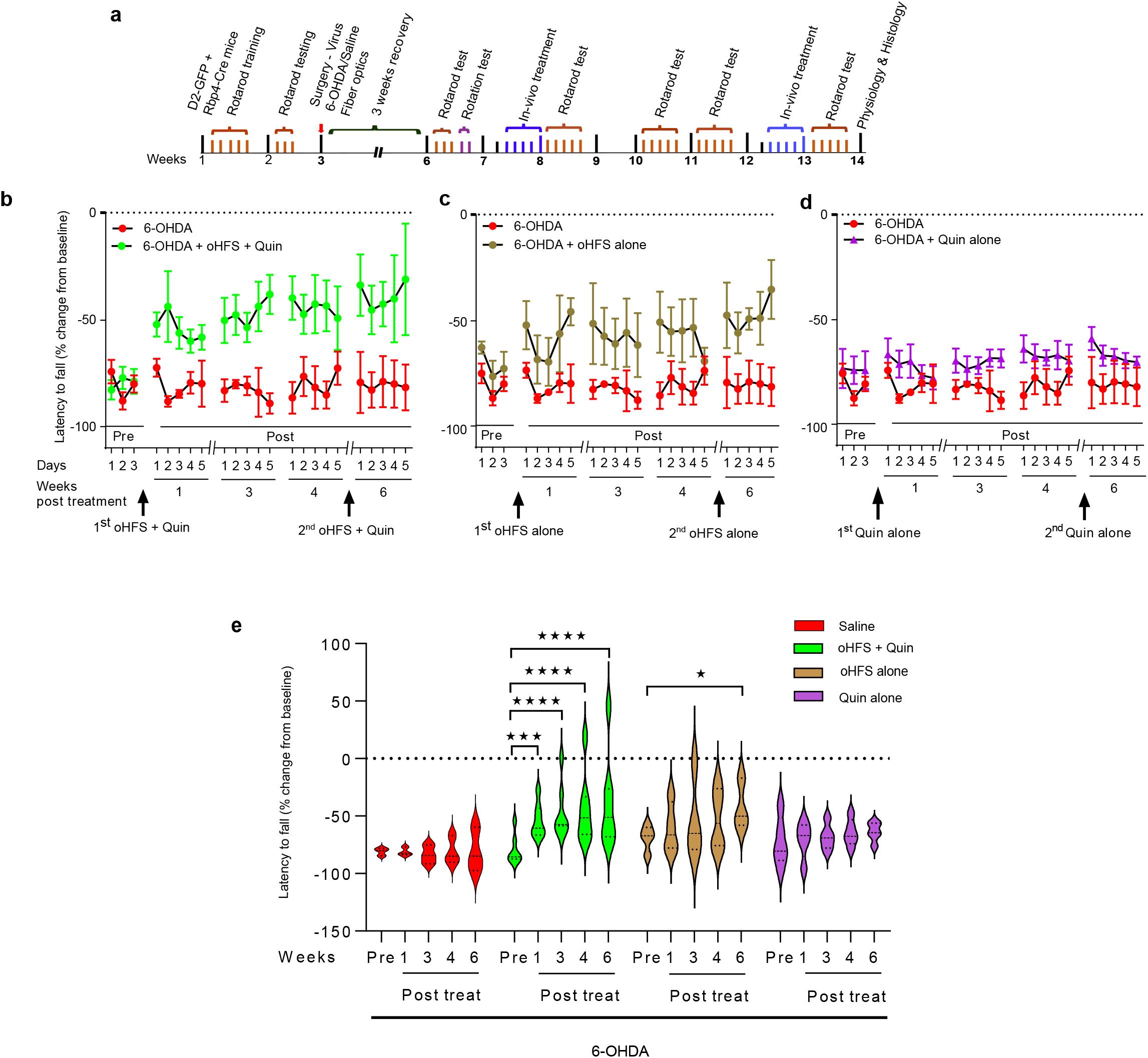
Persistent improvement in rotarod performance with D2 receptor agonist and oHFS in 6-OHDA mice. (a) Experimental timeline for *in-vivo* treatment and rotarod testing in 6-OHDA lesioned mice. (b) Rotarod performance before and after oHFS/Quin treatments in 6-OHDA lesioned mice 5 days/week during weeks 1, 3, and 4 post stimulation with a 2^nd^ oHFS/Quin treatment during week 5, and final testing in week 6 (green, n=7), plotted with untreated controls (red, n=3). (c&d) Rotarod performance before and after oHFS alone in 6-OHDA lesioned mice (brown, n=4), or quinpirole alone (purple, n=5), plotted with untreated controls (red, n=3). (e) Summary graph of rotarod performance in saline (red), oHFS/Quin (green), oHFS alone (brown), and quinpirole alone (purple). Each pretreatment plot represents average of 3 days of testing. Post treatment plots are averages of 5 days of testing. Data are presented as the % change in performance relative to pre-lesioned baseline performance. Statistics: *p ≤ 0.05, ***p ≤ 0.001, ****p < 0.0001. Three-way ANOVA, multiple comparisons with post-hoc Fisher’s LSD test in (e).

We then tested whether the optical stimulation or quinpirole treatment might improve motor performance independently. In one group, oHFS was administered for five sessions, as described above, but without quinpirole injection. We observed only weak improvement in rotarod performance during weeks 1, 3 and 4 post treatment in 6-OHDA lesioned mice (Fig 2c, 2e). A second oHFS treatment between weeks 4 and 6, induced statistically significant improvement in performance relative to pretreatment (Fig 2c, 2e). We next examined whether D2 receptor activation alone could improve motor performance in PD mice. Quinpirole was administered once daily for five days to a 6-OHDA lesioned mice followed by rotarod testing. Quinpirole alone did not improve motor performance in lesioned animals tested during 1, 3 and 4 weeks post treatment (Fig 2d, 2e). Additional quinpirole treatment during the 5^th^ week did not improve their rotarod performance during week 6 (Fig 2d, 2e).

### Combined optogenetic stimulation and quinpirole treatment down-regulates vesicular glutamate transporter 1 expression in DLS of 6-OHDA mice

Type 1 vesicular glutamate transporter (VGLUT1) is highly expressed in cortical pyramidal neurons and its expression correlates with the strength of glutamatergic transmission in dorsal striatum (Du et al., 2020; Fremeau et al., 2001; Wilson et al., 2005; Wojcik et al., 2004). Structural changes associated with LTP induction are associated with increased VGLUT1 expression, while LTD reduces VGLUT1 (Balschun et al., 2010; Nakakubo et al., 2020). Several lines of evidence support the idea that dopamine neuron loss in PD is associated with enhanced glutamatergic activity in dorsal striatum, and an increase in VGLUT1 expression (El Arfani et al., 2015; Kashani et al., 2007). To assess possible changes in striatal VGLUT1 expression in our experimental conditions, we measured VGLUT1 immunostaining fluorescence intensity in DLS of brain slices from 6-OHDA lesioned mice, with and without oHFS/Quin treatment (Fig 3a, 3b). Unilateral 6-OHDA lesions did not significantly affect VGLUT1 immunoreativity in the lesioned hemisphere relative to the non-lesioned side. However, oHFS/Quin treatment of 6-OHDA lesioned mice induced a decrease in VGULT1 expression on the lesioned/optically stimulated side compared to the non-lesioned hemisphere (Fig 3b, 3c). This change in VGLUT1 expression is consistent with an induction of LTD in the stimulated hemisphere and decreased cortical glutamatergic transmission in DLS following in vivo oHFS/Quin treatment.

**Figure 3.**
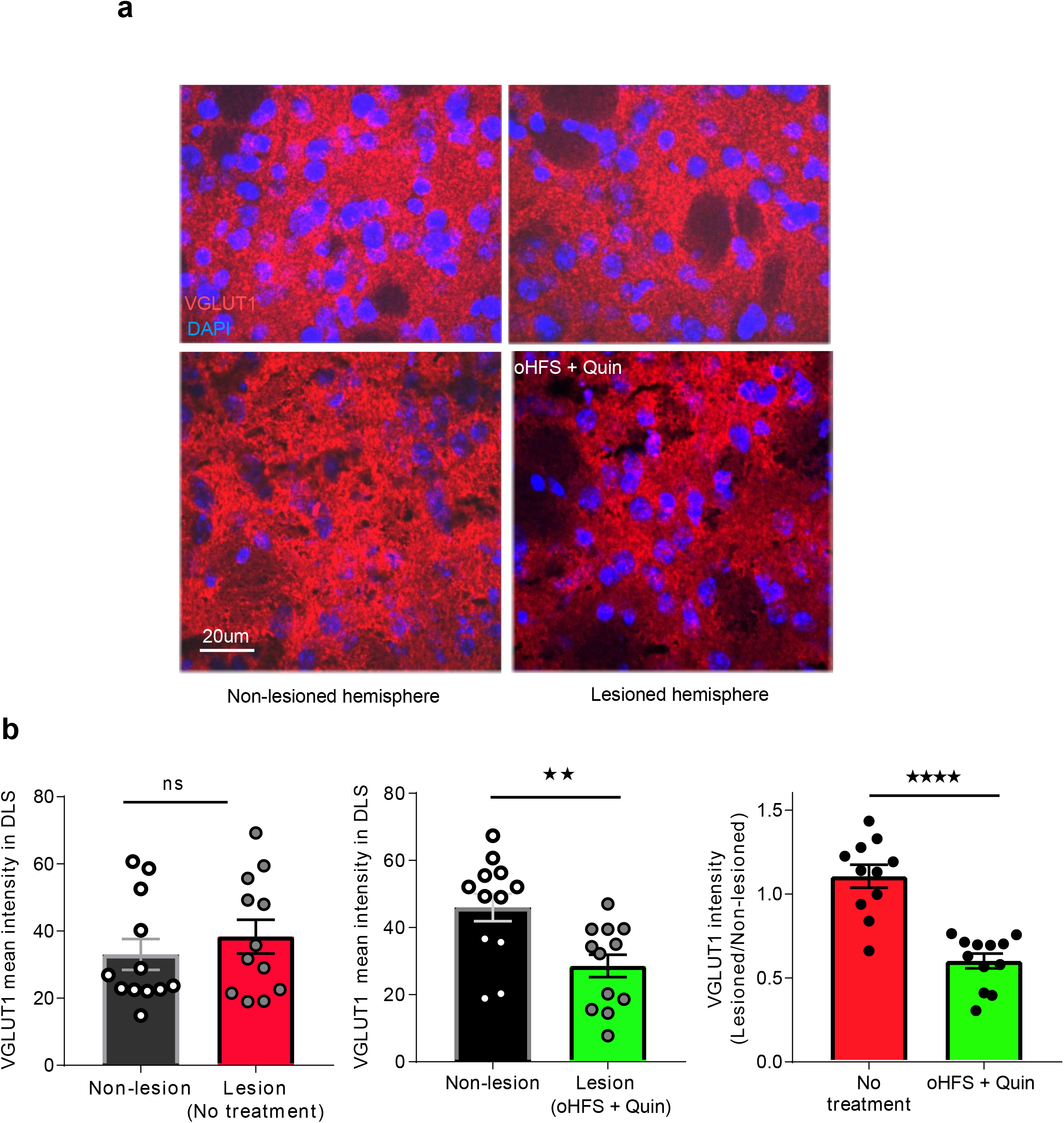
Differential expression of VGLUT1 in 6-OHDA mice that received oHFS/Quin treatment in vivo. (a) Coronal brain section with VGLUT1 immunostaining (red) from unilateral 6-OHDA lesioned mouse. Non-lesioned hemisphere (left panels, top & bottom). Lesioned hemisphere (right panels, top & bottom). No treatment (top right), oHFS/Quin treatment (bottom right). (b) Summary of VGLUT1 intensity on non-lesion side vs lesioned side with no treatment (left, N=12 sections from 3 mice) and non-lesioned vs lesioned with oHFS/Quin treatment (middle, N=12 sections from 3 mice). Ratio of VGLUT1 intensity between lesioned and non-lesioned side in treatment and no treatment groups (right). Scale bar and statistics: 20 μm in (a). **p ≤ 0.005, ****p ≤ 0.0001. Paired t-test, two-tailed in (b, middle), Unpaired t-test, two-tailed in (b, right).

### Combined optical stimulation of cortical inputs with bath application of quinpirole induces LTD in D2-MSNs in brain slices from control and 6-OHDA-lesioned mice

Cortical projections are the principal source of excitatory input to the DLS, and DA differentially modulates the strength of these synapses. DA-induced modulation of cAMP is a critical determinant of the valence and magnitude of plasticity at corticostriatal synapses (Augustin et al., 2014; Calabresi et al., 2007b; Gerfen, 2000; Lerner and Kreitzer, 2011). In D2-MSNs, DA lowers cAMP levels, and these conditions, in combination with mGluR5 activation, and voltage gated calcium channel activity supports the induction of LTD (Lerner and Kreitzer, 2012). In Parkinsonian conditions, loss of DA dysregulates cAMP signaling, leading to increased levels in D2-MSNs, which suppresses LTD induction and promotes LTP of excitatory synapses (Augustin et al., 2014; Calabresi et al., 2007b, 2007b; Rylander et al., 2013). As electrical stimulation can activate many afferent cell types, we tested whether optogenetic activation of M1 cortical inputs to DLS selectively could induce synaptic plasticity in brain slices from our control and 6-OHDA lesioned mice. We recorded from identified D2-MSNs (Fig 1d) in tissue slices from sham surgery control mice and 6-OHDA lesioned animals. For consistency with the behavioral studies, the animals used for electrophysiology were first trained on rotarod for 5 days prior to 6-OHDA treatment (Fig 4a). Optogenetic control of layer 5 M1 projections to striatum was accomplished with Rbp4-dependent ChR2 expression on corticostriatal terminals (see Fig 1a).

**Figure 4.**
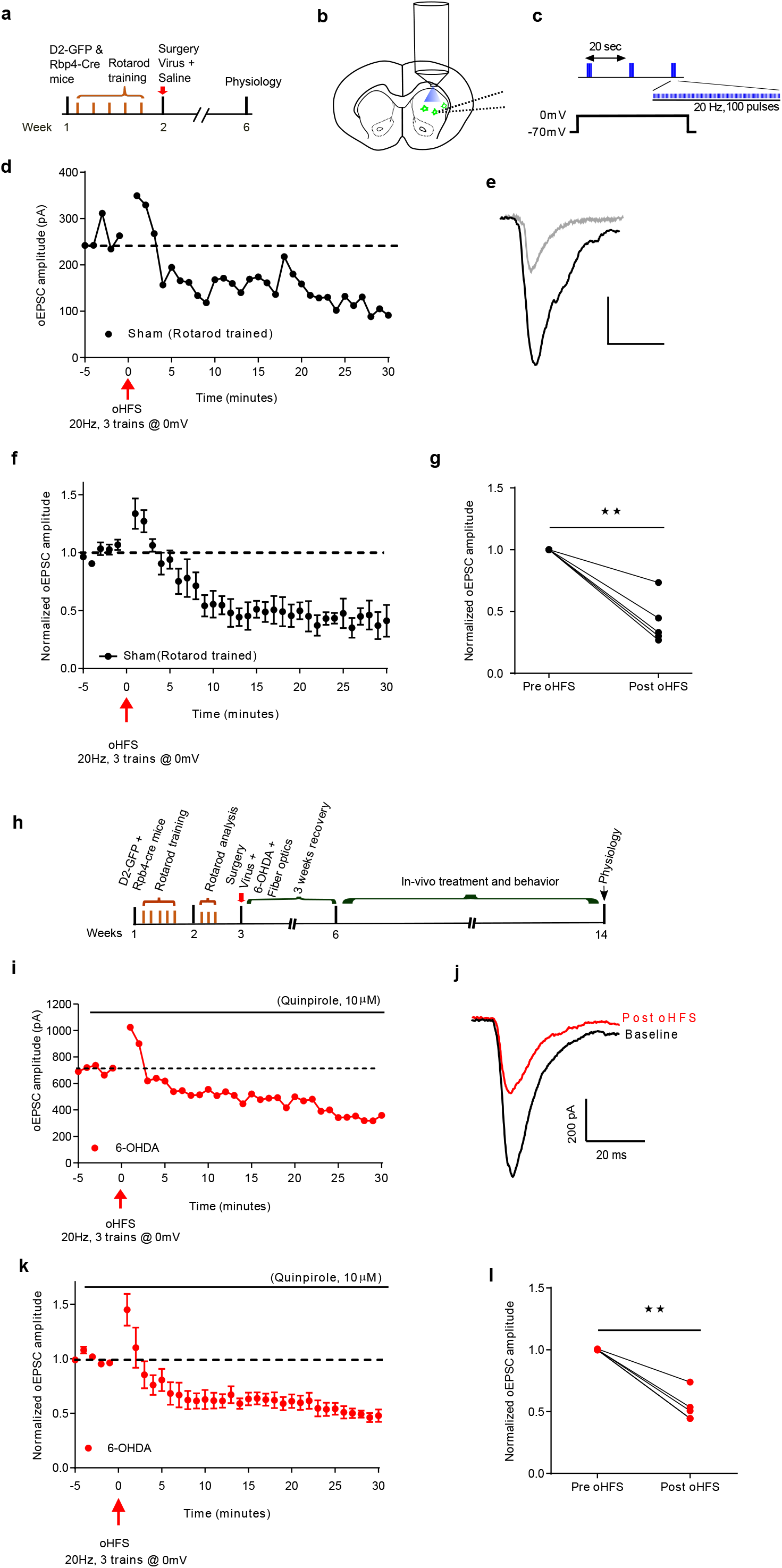
LTD induction with optical high frequency stimulation in brain slices from sham and 6-OHDA lesioned mice. (a) Experimental timeline for ex-vivo slice physiology experiments from control mice. (b) Schematic of recording from D2-GFP neurons and optical stimulation with microscope objective (c) Schematic of oHFS protocol for electrophysiology. While holding the D2-GFP neuron at 0mV 3 trains of 100 blue pulses at 20Hz were delivered with 20 second inter train intervals. (d) Representative example of the oEPSC amplitudes to single optical stimulation during baseline, 5 min prior to oHFS, and after oHFS. (e) Representative trace showing oEPSCs during baseline (black) and 20 min after oHFS (grey). (f) Averaged oEPSC amplitudes normalized to baseline (N=5 cells from 3 mice) before and after oHFS. (g) Average of normalized oEPSCs during baseline and the last 10 min post oHFS for each recorded neuron. (h) Experimental timeline for ex-vivo slice physiology experiments from 6-OHDA lesioned mice. (i) Representative example of the oEPSC amplitudes to single optical stimulation during baseline, 5 min prior to oHFS, and after oHFS in a brain slice from a 6-OHDA lesioned mouse. (j) Representative trace showing oEPSCs during baseline (black) and 20 min after oHFS (red). (k) Averaged oEPSC amplitudes normalized to baseline (N=5 cells from 4 mice) before and after oHFS in brain slices from 6-OHDA lesioned mice. (l) Average of normalized oEPSCs during baseline and the last 10 min post oHFS for each recorded neuron from 6-OHDA lesioned mice. Scale bar and statistics: 200 ms and 200 pA. *p ≤ 0.05, **p ≤ 0.005. Paired t-test in (g & l).

In brain slices from control mice, whole-cell voltage clamp recordings from D2-MSNs yielded robust optically-evoked excitatory post synaptic currents (oEPSCs) from D2-MSNs ipsilateral to the sham injection (Fig 4b). After 5 min of stable baseline recording, oHFS with 3 trains of 20Hz, 100 pulses/train, 20 sec inter-train interval was given, while holding the cell at 0mV (Fig 4c). We monitored oEPSCs for 30 min post oHFS (Fig 4d, 4e), and observed a reduction in oEPSC amplitude relative to baseline in all cells tested (Fig 4f, 4g). In our previous studies, the prevalence of LTD with electrical HFS alone was 67% in slices from control animals (Augustin et al., 2014) but here we observed LTD induction in 100% of the cells tested, reflecting a higher efficacy of plasticity induction with oHFS.

Previous studies in PD mouse models reported a lack of LTD induction using electrical HFS methods in DA depleted slices, and that this form of LTD could be rescued with D2 receptor activation (Kreitzer and Malenka, 2007). In light of these results and our persistent behavioral effects following in vivo oHFS/Quin treatment, we tested the effects of oHFS in combination with bath application of the D2 agonist quinpirole (10 μM) on oEPSC amplitudes in D2 MSNs from 6-OHDA lesioned mice that had undergone behavioral assessments (Fig 4h). Baseline oEPSCs were measured in the presence of quinpirole for 5 min, and oHFS was given to induce oLTD. We observed significant reduction of oEPSC amplitudes in all recorded cells (Fig 3 i, j, k,l).

We next tested whether *in-vivo* oHFS/Quin treatment could occlude in vitro LTD induction with oHFS and quinpirole. Brain slices were collected from 6-OHDA lesioned mice that had undergone oHFS/Quin treatment 24 hrs before. Recording from D2-MSNs in these slices, we found that the combination of oHFS with quinpirole induced robust LTD that was not different from that obtained in tissue slices from sham controls or untreated 6-OHDA lesioned mice. Despite the in vivo treatment, we did not see evidence of LTD occlusion.

### Increased intrinsic excitability of D2-MSNs in tissue slices from 6-OHDA lesioned mice following *in vivo* LTD induction

DA depletion is expected to increase excitability of D2-MSNs due reduced D2 receptor activity. Published observations of changes in D2-MSN intrinsic excitability in PD models are mixed, with some studies reporting increase excitability, and others showing decreased excitability, or no effect (Azdad et al., 2009; Fieblinger et al., 2014; Ketzef et al., 2017). This discrepancy could be due to the timing of experiments after DA depletion, or differences in lesion protocols. Here we tested the effects of DA depletion and *in-vivo* oHFS/Quin treatment on D2-MSN excitability in tissue slices from our 6-OHDA lesioned animals. In whole-cell current clamp recordings, we monitored firing rates in response to a series of depolarizing current injections. In recordings from D2-MSNs in tissue slices from 6-OHDA-lesioned mice, we did not see a significant difference in firing rate or spike-frequency adaptation relative to slices from sham control mice. Interestingly, firing rates were higher in tissue slices from oHFS/Quin treated mice compared to 6-OHDA lesioned animals without treatment (Fig 5a, 5b). When we measured threshold current, we found a significant shift to a lower threshold in the neurons from oHFS/Quin treated mice compared to 6-OHDA lesioned animals (Fig 5c). These findings suggest that the oHFS/Quin treatment leads to a homeostatic shift in excitability of D2-MSNs, reflecting the weaker excitatory drive to these neurons due to LTD induction.

**Figure 5.**
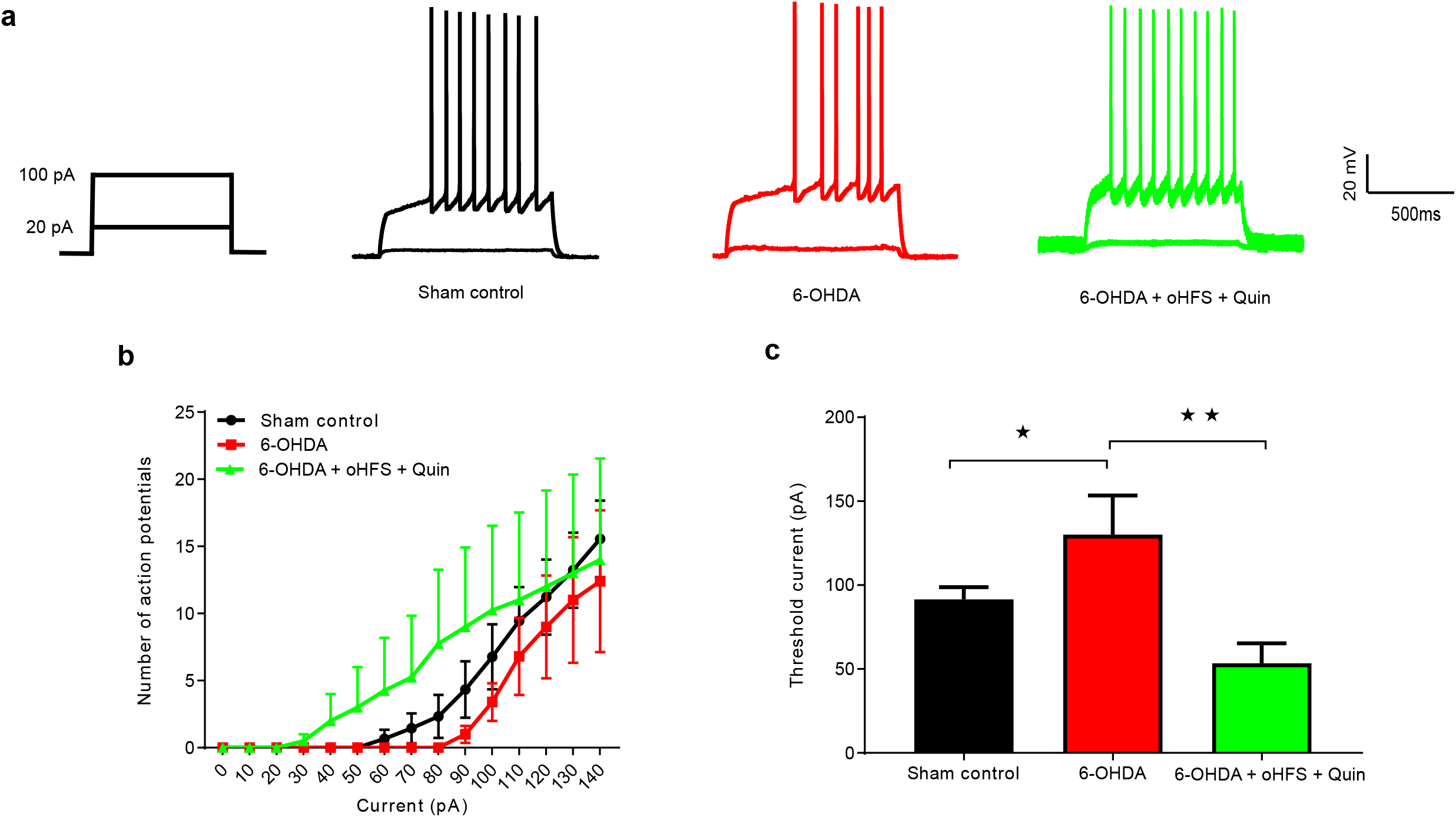
Action potential threshold changes following oHFS/Quin treatment in 6-OHDA lesioned mice. (a) Schematic of step currents (right). Representative traces of action potentials to 20 and 100pA step currents in D2-GFP neurons from sham control (black), 6-OHDA (red), and 6-OHDA that received oHFS/Quin treatment (green). Treatment timeline for these animals is illustrated in Fig 4h. (b) Number of action potentials evoked by injected current for sham control (black), 6-OHDA lesioned (red), and oHFS/Quin treated (green) groups. (c) Threshold current required to elicit action potential in sham (N=7), 6-OHDA (N=10), and 6-OHDA + oHFS/Quin stimulation (n=4). Scale bar and statistics: 500 ms and 20 mV. *p ≤ 0.05, **p ≤ 0.005. Ordinary one-way ANOVA with Fischer’s LSD post-hoc test (c).

## Discussion

Death of dopamine neurons in PD patients leads to progressively worsening motor deficits, which include tremor, rigidity, difficulty initiating movement, loss of coordination, and a dramatic decline in motor learning. Dopamine replacement therapy is the gold standard frontline treatment, with remarkable efficacy in reducing symptoms and restoring motor control. Unfortunately, over time, decreased therapeutic efficacy, combined with the development of uncontrolled movements or dyskinesia, along with psychiatric side effects limit this treatment strategy (Barbeau, 1976; Lane, 2019; Marsden, 1994; Mosharov et al., 2015; Thanvi and Lo, 2004). Alternative pharmacological approaches that target cholinergic or adenosine receptor systems do not relieve symptoms to the same degree. Deep brain stimulation (DBS) is effective for many patients, but this method is not appropriate for all patients, and there are issues with the long-term effectiveness of the implanted electrodes due to impedance changes (Nassery et al., 2016; Okun, 2012; Satzer et al., 2020; Voon et al., 2013). Other commonly reported side effects include impairment of axial motor performance and cognition, along with mood effects including apathy, anxiety, impulsiveness, and depressive symptoms (Hartmann et al., 2019). While it is important to recognize the remarkable relief that DA replacement brings to PD patients, the limits of these treatments emphasizes the need to identify novel approaches to augment or replace existing therapies.

Our previous studies demonstrated that D2 receptor blockade combined with a rotarod motor task induces aberrant motor learning in mice, which persists for nearly two weeks. The persistence of this aberrant motor learning is thought to be dependent upon synaptic plasticity mechanisms, principally within D2 pathway (Beeler et al., 2012). Those studies also demonstrated that pairing of D2 blockade with the motor activity is critical for aberrant learning, as the same D2 blocker treatment administered in the home cage had no effect on motor learning or rotarod performance (Beeler et al., 2012). In human PD patients, motor tasks conducted in the absence of L-DOPA impedes future performance, supporting the idea that aberrant learning occurs in a task-specific manner in low dopamine conditions (Anderson et al., 2014). Interestingly, tasks learned in the presence of L-DOPA are performed more rapidly and with higher accuracy in subsequent trials. These data indicate that DA contributes to the plasticity required for learning and maintenance of optimal motor performance.

Endocannabinoid mediated LTD is a well-characterized form of synaptic plasticity in DLS. Activation of D2 receptors, along with L-type calcium channels, and mGluR5 on D2-MSNs supports the production and release of endocannabinoids (eCB) (Adermark and Lovinger, 2007; Mathur and Lovinger, 2012). While the mechanisms controlling eCB release are not known (Gerdeman et al., 2002) their activation of CB1 receptors present on the pre-synaptic glutamatergic terminals is known to decrease glutamate release and suppress excitatory drive. In DA depleted conditions, eCB-mediated presynaptic inhibition is reduced, leading to enhanced glutamate release and greater excitability in the D2 pathway. *Ex-vivo* slice physiology studies have demonstrated that DA depletion impairs LTD induction at D2-MSNs, but this plasticity can be rescued with D2 receptor activation (Kreitzer and Malenka, 2007). Here we explored the in vivo behavioral consequences of exploiting this stimulation strategy using optogenetic activation of corticostriatal projections.

Optogenetic methods have provided powerful tools for assessing the physiological and behavioral impact of specific neuronal population or connections. Here we have used those methods to induce synaptic plasticity in the motor cortical projections to striatum, with the goal of relieving the motor deficits due to the loss of nigrostriatal DA. Our results show that a stimulation paradigm that reliably induces LTD in brain slices results in marked improvement in rotarod performance. Optical high frequency stimulation of corticostriatal projections, in combination with D2 receptor agonist administration for 5 days, results in motor improvement that persists for over 4 weeks. Neither optical stimulation nor quinpirole treatments alone improved motor performance to the same degree as the combined treatment, suggesting that both were necessary to induce plasticity at corticostriatal synapses. In tissue slices from 6-OHDA lesioned mice that had undergone the combined stimulation paradigm, we observed a decrease in VGLUT1 expression in the stimulated hemisphere relative to the unstimulated side. This change in VGLUT1 levels is consistent with the induction of LTD of excitatory inputs to that region. Our physiological results in brain slices show that D2 receptor activation and high frequency optical stimulation induces LTD of the cortical projections to D2-MSNs in DLS. Thus, our combined treatment paradigm is likely inducing a decrease in excitatory drive to D2-MSNs in vivo, leading to long-lasting motor improvement.

DA levels in DLS are important determinants of corticostriatal synaptic strength, but DA also modulates the intrinsic excitability of D2-MSNs. Several studies have demonstrated reduced excitability of D2-MSNs in Parkinsonian conditions, which likely reflects homeostatic downregulation of firing activity in response to increased excitatory drive to these neurons. Here we tested whether D2-MSN excitability is altered in 6-OHDA animals and the effects of in vivo combined stimulation in these mice. In line with published reports, we observed reduced firing rates in D2-MSNs of 6-OHDA animals. After in vivo LTD induction, we found increased firing rates and hyperpolarized action potential thresholds in these neurons, supporting a homeostatic shift in the intrinsic D2-MSN excitability levels. Together, these observations are consistent with the induction of LTD and decreased excitatory drive to DLS following in vivo oHFS/Quin treatment.

In our *ex-vivo* slice physiology experiments, we looked for evidence of occlusion of LTD in tissue from mice that had experienced oHFS/Quin treatment in vivo. Interestingly, the LTD magnitude and prevalence were similar to that seen in slices from 6-OHDA lesioned mice that had not received in vivo treatment. This could be due to the lack of specificity in choosing the cells that have already undergone LTD. Another possibility is that despite the strong stimulation in vivo, LTD was not saturated in the synapses of the recorded neurons.

The observed improvement in motor function following in vivo oHFS/Quin treatment is remarkable in its persistence for multiple weeks. Based upon our previous studies, we expected that stimulation-induced motor improvement would decline to post-lesion levels with repeated testing, due to the low DA conditions in our mice (Beeler et al., 2012). The persistent motor improvement implies that our oHFS/Quin paradigm is leading to stable changes in synaptic strength, in a manner that is fundamentally different, possibly due to enhanced Ca^2+^ entry through ChR2 channels presynaptically (Lerner et al., 2016). The requirement of co-stimulation of D2 receptors highlights the key contribution of postsynaptic signaling mechanisms. Further study will be needed to identify the cellular mechanisms that mediate the changes in physiology that support the remarkable persistence of the rescue of motor performance in these animals.

The in vivo oHFS/Quin treatment resulted in improved motor performance, but not to baseline levels, even after a second treatment (Fig 2). Less than complete recovery could result from the limited specificity of our stimulation paradigm. Our use of an Rbp4-cre dependent expression system selectively induces ChR2 in layer 5 pyramidal neurons, which resulted in reliable presynaptic activation of M1 cortical inputs to DLS. However, we are not likely activating *all* rotarod-related inputs, which could limit the extent of recovery. Alternatively, inducing LTD in DLS neurons that are *not* involved in rotarod performance may limit the efficacy of these methods. In addition, cortical inputs to the D1 pathway are active in our oHFS stimulation, and plasticity in these connections could also compromise motor improvement. Future testing of engram-specific stimulation could improve the performance levels following in vivo treatments of this type (Josselyn and Susumu, 2020).

Our results suggest an alternative therapeutic strategy for induction of long-lasting relief of motor symptoms in Parkinson’s disease. Previous human studies have demonstrated the safety, tolerability, and potential efficacy of AAV vectors for gene therapy in the putamen of PD patients (Muramatsu et al., 2010). In addition, the use of optogenetics is already underway in clinical trials for human retinitis pigmentosa, but its application to neurological disease is in its infancy. The persistence of the relief of motor symptoms in 6-OHDA lesioned mice in our studies suggests that optogentic induction of synaptic plasticity at cortical inputs to DLS may prove to be an efficacious alternative or adjuvant to available treatments for relieving Parkinsonian motor symptoms.

## Acknowledgements

This work was supported by NIH grants NS095374 and NS110371

## Author contributions

CA designed experiments, and contributed to all the experimental approaches, generated all the figures, wrote the first draft of the manuscript and incorporated suggested edits. JT and BR carried out surgical procedures, conducted motor performance testing, provided feedback on figures and helped edit the manuscript. XZ and DSM provided funding for the project, designed the experiments, consulted with CA, JT, and BR during the experiments, edited the figures and the manuscript.

## STAR* Methods

Key Resources Table

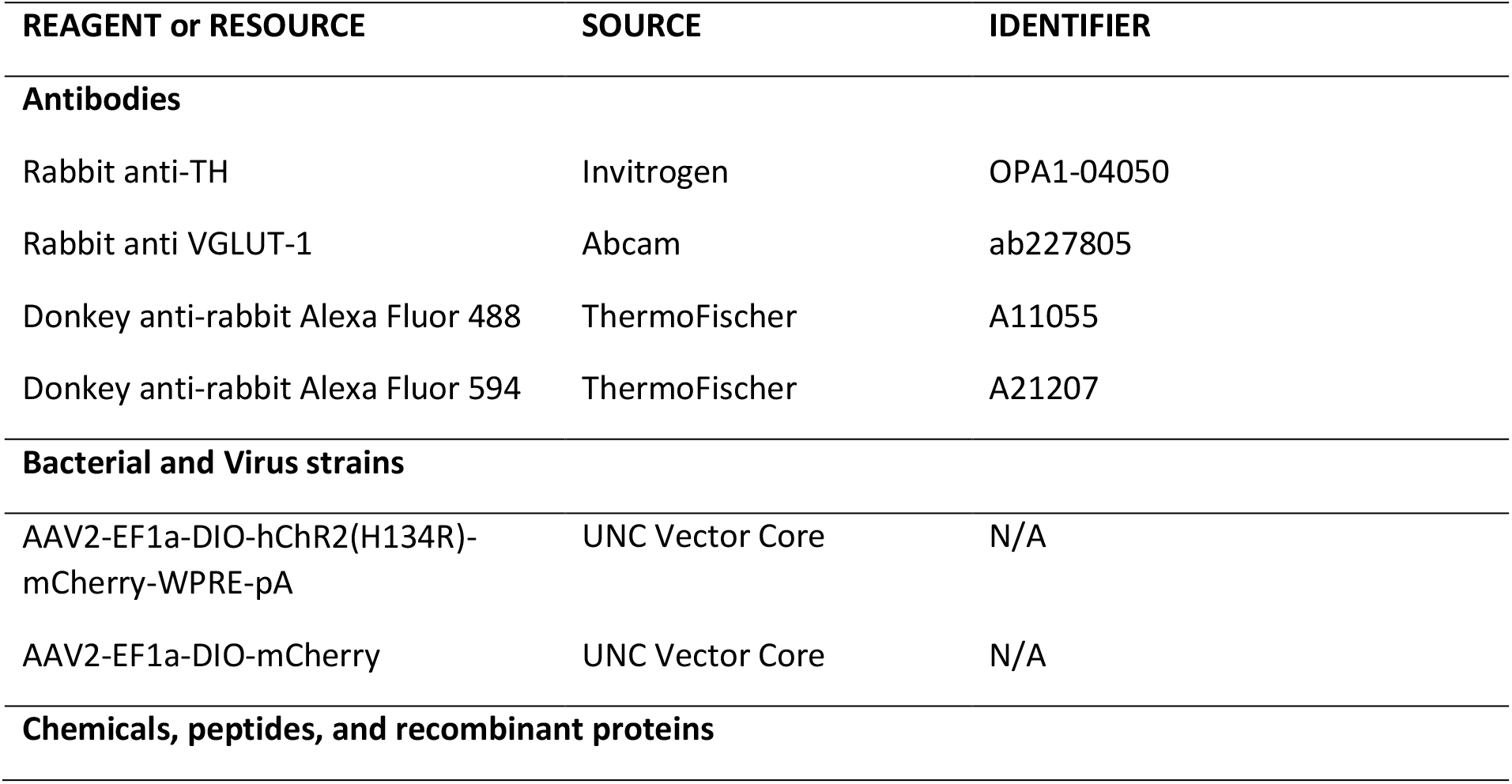
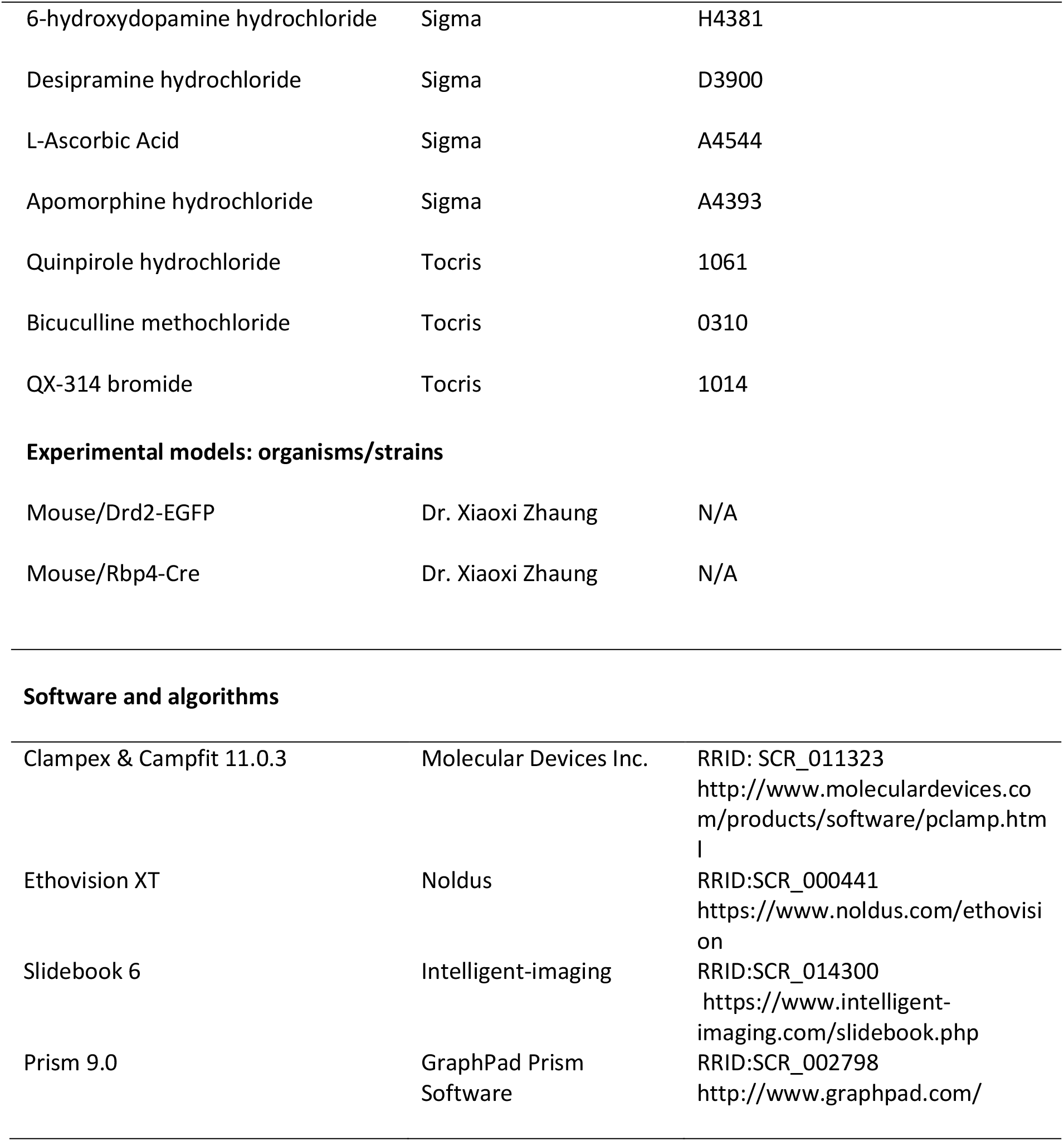

| REAGENT or RESOURCE | SOURCE | IDENTIFIER |
| --- | --- | --- |
| **Antibodies** |  |  |
| Rabbit anti-TH | Invitrogen | OPA1-04050 |
| Rabbit anti VGLUT-1 | Abcam | ab227805 |
| Donkey anti-rabbit Alexa Fluor 488 | ThermoFischer | A11055 |
| Donkey anti-rabbit Alexa Fluor 594 | ThermoFischer | A21207 |
| **Bacterial and Virus strains** |  |  |
| AAV2-EF1a-DIO-hChr2(H134R)-mCherry-WPRE-pA | UNC Vector Core | N/A |
| AAV2-EF1a-DIO-mCherry | UNC Vector Core | N/A |
| **Chemicals, peptides, and recombinant proteins** |  |  |
| 6-hydroxydopamine hydrochloride | Sigma | H4381 |
| Desipramine hydrochloride | Sigma | D3900 |
| L-Ascorbic Acid | Sigma | A4544 |
| Apomorphine hydrochloride | Sigma | A4393 |
| Quinpirole hydrochloride | Tocris | 1061 |
| Bicuculline methochloride | Tocris | 0310 |
| QX-314 bromide | Tocris | 1014 |
| **Experimental models: organisms/strains** |  |  |
| Mouse/Drd2-EGFP | Dr. Xiaoxi Zhaung | N/A |
| Mouse/Rbp4-Cre | Dr. Xiaoxi Zhaung | N/A |
| **Software and algorithms** |  |  |
| Clampex & Clampfit 11.0.3 | Molecular Devices Inc. | RRID: SCR_011323<br>http://www.moleculardevices.com/products/software/pclamp.html |
| Ethovision XT | Noldus | RRID:SCR_000441<br>https://www.noldus.com/ethovision |
| Slidebook 6 | Intelligent-imaging | RRID:SCR_014300<br>https://www.intelligent-imaging.com/slidebook.php |
| Prism 9.0 | GraphPad Prism Software | RRID:SCR_002798<br>http://www.graphpad.com/ |

### Animals

All animal procedures were performed according to the approved protocol of the Institutional Animal Care and Use Committee (IACUC) at the University of Chicago. To generate D2-GFP and Rbp-4 Cre double transgenic mice, Drd2-eGFP (D2-GFP) BAC transgenic mice on C57BL/6 background were crossed with Rbp-4 cre mice. All the experiments were conducted on adult (>8 weeks; both male and female) hemizygous mice. Animals were maintained on 12-hour light/dark cycle. All behaviors were conducted during daytime.

### Stereotaxic surgeries and post-surgery care

Stereotaxic surgeries were performed on hemizygous adult Drd2-eGFP and Rbp4-Cre double transgenic mice. All the mice were trained on rotarod prior to the surgeries. Mice were maintained on isoflurane anesthesia throughout the surgery. Mouse heads were fixed on a stereotaxic frame (Kopf instruments). Desipramine (25mg/kg) was injected intraperitoneally to block the norepinephrine/5HT uptake 30 min prior to the 6-hydroxy dopamine (6-OHDA) injections. All the injections were made relative to Bregma. To deplete dopaminergic neurons in SNc, 1μl of 6-OHDA (4μg/μl) in saline containing 0.02% ascorbic acid was unilaterally injected into MFB (coordinates: AP −1.05mm, ML +1.2mm, DV −1.2mm) using a microinjection system (WPI, UMP3T-1) at a rate of 120nl/min. One cohort of mice was administered 6-OHDA injections in DLS (coordinates: AP −0.18mm, ML ±2.4mm, DV −3.0mm; Fig S1). The 6-OHDA solution prepared fresh daily. As the 6-OHDA injections cause severe weight loss, animals were closely monitored for their health and supplemented with diet gel (ClearH20) and saline injections until body weights recovered. After surgery, animals were allowed 3 weeks for recovery before behavioral testing. For sham controls, 1μl saline with 0.02% ascorbic acid was injected using the same coordinates. Mice were kept on thermal pad throughout the surgery to avoid hypothermia. To express channel rhodopsin in layer V of motor cortex, 500nl of AAV2-DIO-hChR2 (H134R)-mCherry was injected unilaterally (coordinates: AP −0.1 mm; ML +1.5 mm; DV −1.2 mm). The injection canula was left in place for about 10 min to allow spread of virus into the tissue. For in-vivo optogenetic stimulation of layer V motor cortical inputs in DLS, fiber optic cannulae were implanted (coordinates: AP +0.80mm, ML ±2.2mm, DV −2.7mm).

### Behavior

Animals that failed to recover normal body weight within 3 weeks were excluded from behavioral testing. To assess the unilateral lesion-induced motor behaviors, and identify experimental subjects, mice were tested using an apomophine challenge rotation test, rotarod motor testing, and post hoc TH immunocytochemistry (see below). The rotation test was conducted on two consecutive days. Animals were habituated to the behavioral room for 60 min prior to the testing. On day 1, mice were injected subcutaneously with saline and placed in white plastic cylinders (22.1 cm diameter) for 30 min and then returned to their home cages. On the second day, same test was repeated with subcutaneous injection of apomorphine (0.5 mg/kg s.c.) and the behavior was recorded with live video streaming for 30 min. Full body turning (360°) was counted as a rotation. The number of contra- and ipsilateral rotations (relative to the 6-OHDA lesioned hemisphere) were measured using post hoc video analysis with Ethovision software. Data were expressed as apomorphine induced net contralateral rotations. The threshold for inclusion of experimental subjects required >20 net contralateral rotations after apomorphine injection.

To assess the motor coordination, we used rotarod testing, where the mice were placed on an accelerating rotarod (4-40rpm, acceleration: 0.1 rpm/sec) and monitored for the latency to fall. Prior to 6-OHDA lesion, mice were trained on the rotarod with 5 trials/day with 10 min inter-trial interval, for 5 consecutive days. After 3 weeks of recovery from 6-OHDA surgery, mice were tested again on rotarod for 3 days. Data were expressed as change in latency to fall relative to baseline, which is the average of the last three days of training. The threshold for inclusion of experimental subjects was >30% change relative to baseline after 6-OHDA lesion.

The oHFS/Quin stimulation paradigm consisted of 60 min habituation followed by quinpirole administration (5 mg/kg, i.p.), and 30 min later, 5 trains of 20 Hz optical stimulation, 5 min each, with 1 min inter-train interval. Starting the next day after the final in vivo treatment, mice were tested on the rotarod daily to assess motor performance.

### Brain slice preparation

Mice were anesthetized with isoflurane and decapitated. Brains harvested and coronal slices (250 μm) containing DLS were cut using a vibratome (VT 1200S, Leica Microscystems) in ice-cold NMDG solution containing: in mM, 92 N-methyl-D-Glucamine, 2.5 KCl, 1.25 NaH_2_PO_4_, 30 NaHCO_3_, 20HEPES, 25 glucose, 2 thiourea, 5 Na-ascorbate, 3 Na-pyruvate, 0.5 CaCl_2_.4H_2_0, and 10 MgSO_4_.7H_2_0, pH 7.3 to 7.4, bubbled continuously with 95% O_2_ and 5% CO_2_. Slices were incubated for 10 min at 32°C in NMDG solution bubbled with 95% O_2_ and 5%CO_2_, then transferred to HEPES solution containing: in mM, 119 NaCl, 2.5 KCl, 1.25 NaH_2_PO_4_, 30 NaHCO_3_, 20HEPES, 25 glucose, 2 thiourea, 5 Na-ascorbate, 3 Na-pyruvate, 2 CaCl_2_.4H_2_0, and 2 MgSO4.7H_2_0, pH 7.4, bubbled continuously with 95% O_2_ and 5%CO_2_ and perfused at a rate of 20ml/min. Slices were incubated for minimum 30 min in HEPES holding solution at room temperature before electrophysiological recordings. Slices were then transferred into a recording chamber and constantly superfused with aCSF containing: in mM, 125 NaCl, 25 NaHC0_3_, 20 glucose, 2.5 KCl, 2.5 CaCl_2_, 1 MgCl_2_, 1 NaH_2_PO_4_, pH 7.4, bubbled with 95%CO_2_ and 5%O_2_. All the electrophysiological recordings were conducted at room temperature. Medium spiny neurons with GFP reporter were visualized with blue light through water immersion objective (40x) on upright microscope (Zeiss, Axio).

### Electrophysiology

Whole cell patch-clamp recordings were made on D2-GFP neurons in DLS using Multiclamp 200B amplifier (Molecular Devices). Patch pipettes with resistance of 3-5M were pulled using a pipette puller (model P-97, Sutter Instrument, Novato, CA). All the signals were filtered at 2kHz and digitized at 5kHz using Digidata 1550B (Molecular Devices). For whole cell voltage clamp recordings, recording pipette was filled with internal solution containing: in mM, 117 Cs-gluconate, 20 HEPES, 0.4 EGTA, 2.8 NaCl, 20 Glucose, 2.5 ATP, 0.25 GTP, 5 TEA, 5 QX-314, pH 7.3-7.4 with CsOH. All the recordings were conducted with bath application of GABAA anatagonist, bicuculline (20 μM) and the holding potential was −70mV. For whole cell current clamp recordings, internal solution containing: in mM, 145 K-gluconate, 0.1 CaCl_2_.2H_2_0, 5 KCl, 10 HEPES, 0.1 EGTA, 4 Mg-ATP, 0.3 Na-GTP, 4 MgCl_2_, 10 phospho creatinine (disodium salt), pH 7.3-7.4 with KOH. Series resistance was monitored throughout the recording by a hyperpolarizing step (10mV, 1000ms). Data were excluded if series resistance changed >20%.

Optical evoked EPSCs were measured by stimulation of ChR2 expressed on layer V cortical inputs in DLS using blue light pulses (5 ms, 470 nm). For induction f LTD, high frequency stimulation (HFS) consists of three trains of 20 Hz optical stimulation with a 20 second inter train interval, and each train with 100 light pulses was delivered while holding the cell at 0mV. LTD induction protocol was given after obtaining a stable baseline at least for five min. To measure HFS induced LTD, oEPSCs were monitored for 30 min post LTD induction and average oEPSC amplitudes during baseline were compared with the average oEPSC amplitude from the last 10 min recording.

### Histology

Histology for tyrosine hydroxylase (TH) was performed on tissue slices after electrophysiological recording (250 μm). These sections were immersion fixed overnight in 4% paraformaldehyde (PFA) in phosphate buffered solution (PBS), pH 7.4. VGLUT1 immunohistochemistry was conducted using 40 μm cryostat tissue sections (Leica Biosystems) from mice that were anesthetized and transcardially-perfused with 4% PFA in PBS. Brains were harvested and immersed in 4% PFA overnight at 4^0^C and then dehydrated in 30% sucrose in PBS at 4^0^C for 24 hours. For both VGLUT1 and TH immunostaining, slices were washed with PBS three times and then blocked and permeabilized with PBS containing 1% BSA, 10% normal donkey serum and 0.1% Triton X-100 for 2 hours at room temperature followed by primary antibody incubation for 24 hours at 4^0^C. TH immunostaining we used a rabbit polyclonal tyrosine hydroxylase antibody (1:500 dilution, Invitrogen OPA1-04050) and VGLUT1 we used a rabbit monoclonal VGLUT1 antibody (1:500, Abcam ab227805). Slices were washed with 1x PBS, three times and then incubated with the secondary antibody donkey anti rabbit Alexa Fluor 488 or 594 (1:1000 dilution, ThermoFischer: A11055, A21207) for 2 hours at room temperature and washed with 1x PBS. Images were acquired using a Zeiss confocal fluorescent microscope. Post hoc assessment of the 6-OHDA lesion involved comparison of the fluorescent immunostaining intensity on the lesioned relative to the unlesioned hemisphere. Animals with TH staining intensity difference of less than 20% between the hemispheres were excluded from the analysis. VGLUT1 fluorescence intensity was compared between the lesioned and unlesioned hemispheres of 6-OHDA lesioned mice that received oHFS/Quin treatment or no treatment controls.

### Quantification and Statistical analysis

Electrophysiology data was analyzed using Clampex 11 and Clampfit 11 software (Molecular Devices Inc.). Fluorescence intensity was quantified using Fiji (Image J) software. Rotation data was analyzed with Ethovision XT software. All the statistical analyses were conducted using GraphPad Prism 9 (GraphPad Software Inc., CA, USA). Data were shown as mean ± SEM except Fig 2e where data were violin plotted with a center dashed line as median and two dotted lines represent interquartile range. The shape of the plot represents distribution of rotarod data, with wider sections of the plot indicating higher probability. Lower case ‘n’ indicates number of animals and upper case ‘N’ indicates number of cells used in each experiment. Comparisons between two groups used parametric paired t-test or nonparametric Mann-Whitney test, and multiple comparisons with more than two groups were made with ordinary one-way ANOVA or three-way repeated-measures ANOVA with Fisher’s LSD post-hoc test.

---

**Supplementary Figure 1:**
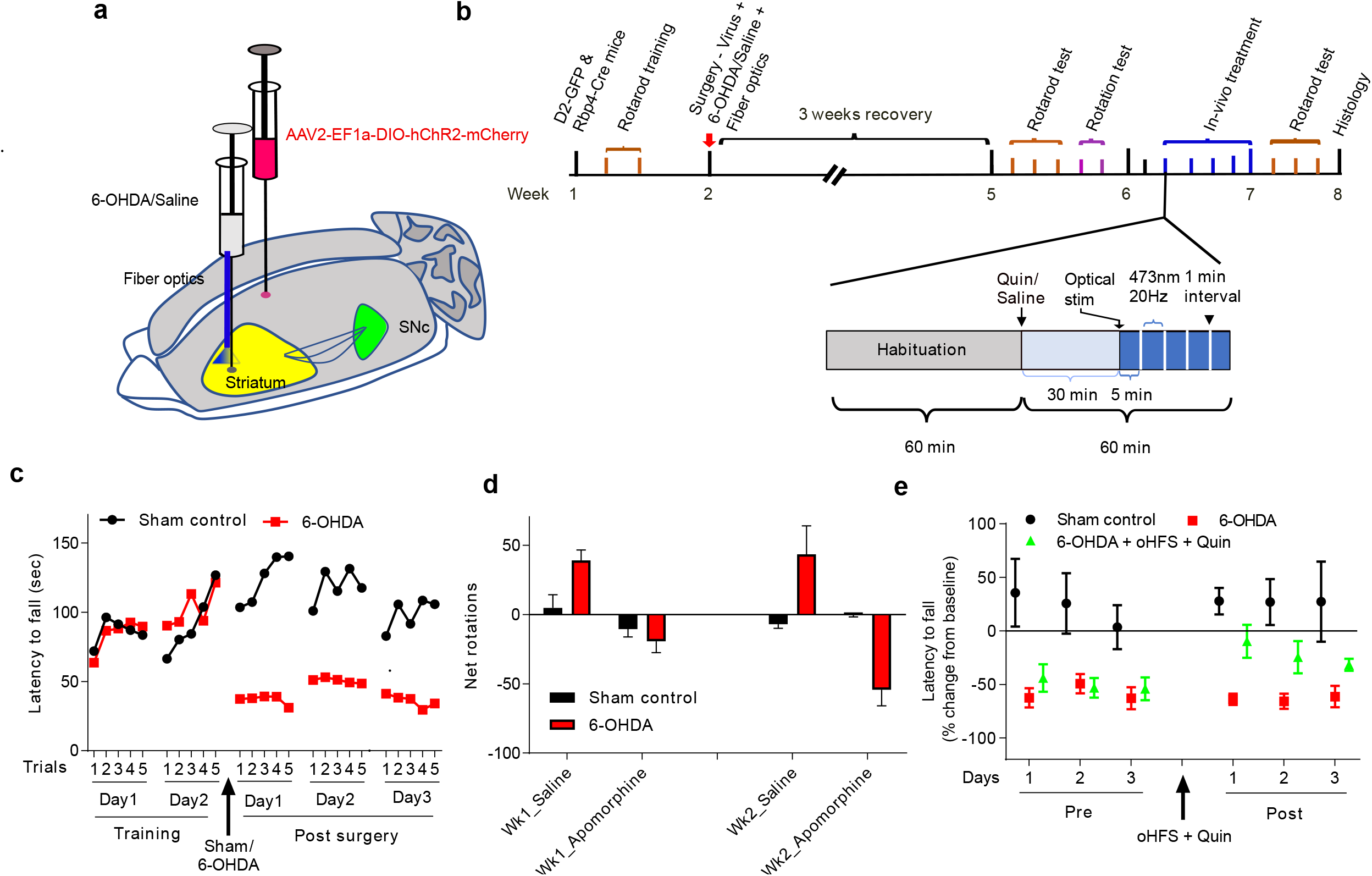
In-vivo oHFS/Quin treatment improves motor performance in mice that received 6-OHDA injections to DLS. a. Schematic of virus administration in M1 cortex, 6-OHDA injection in dorsolateral striatum (DLS) and fiberoptic implantation in DLS. b. Timeline of surgeries, behavioral tests and in-vivo treatment protocol c. Rotarod performance in sham and 6-OHDA lesioned mice during training and 3 weeks after surgery. 6-OHDA mice (red squares, n=8) and sham controls (black circles, n=4). d. Difference in ipsi vs contra lateral rotations with saline and apomorphine 1 and 2 weeks after 6-OHDA/Sham surgeries. e. Rotarod test before and after *in-vivo* oHFS/Quin treatment in 6-OHDA lesioned mice. Treatment group (green triangles, n=3), saline treated 6-OHDA lesioned mice (red squares, n=5), and sham controls (black circles, n=4).

**Supplementary Figure 2:**
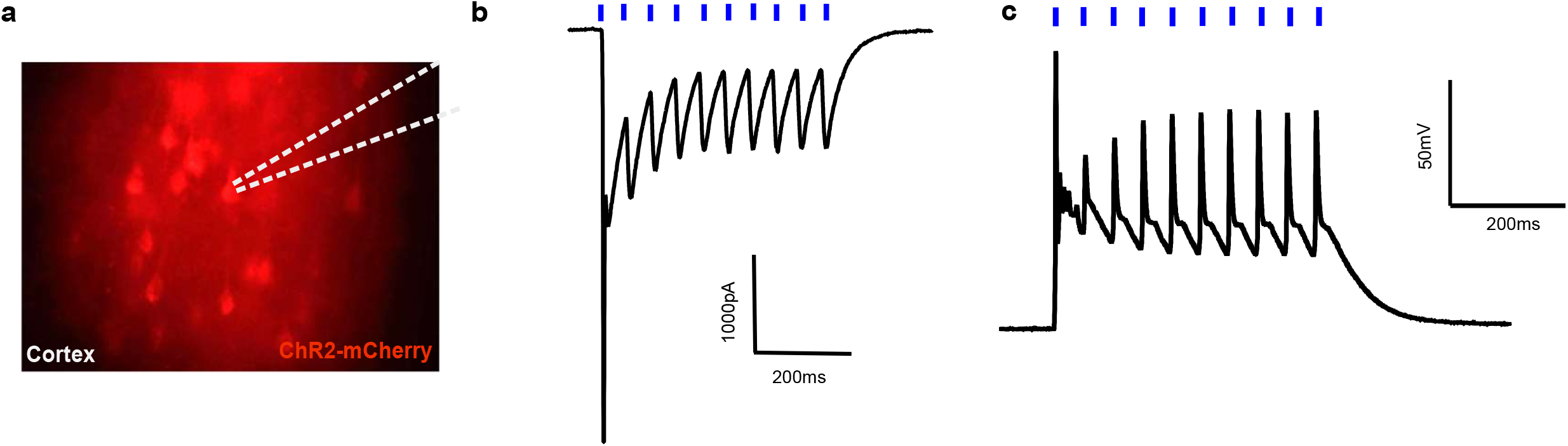
Ex-vivo verification of ChR2 function in motor cortex. a. Coronal brain slice shows expression of mCherry in layer V of motor cortical neurons. b. Representative traces showing inward currents in response to 470 nm light pulses at 20Hz in voltage clamp recording from mCherry expressing cortical neurons. c. Representative trace showing current clamp recording from an mCherry expressing neuron. Each blue light pulse at 20Hz elicit action potential in these neurons.

**Supplementary Figure 3:**
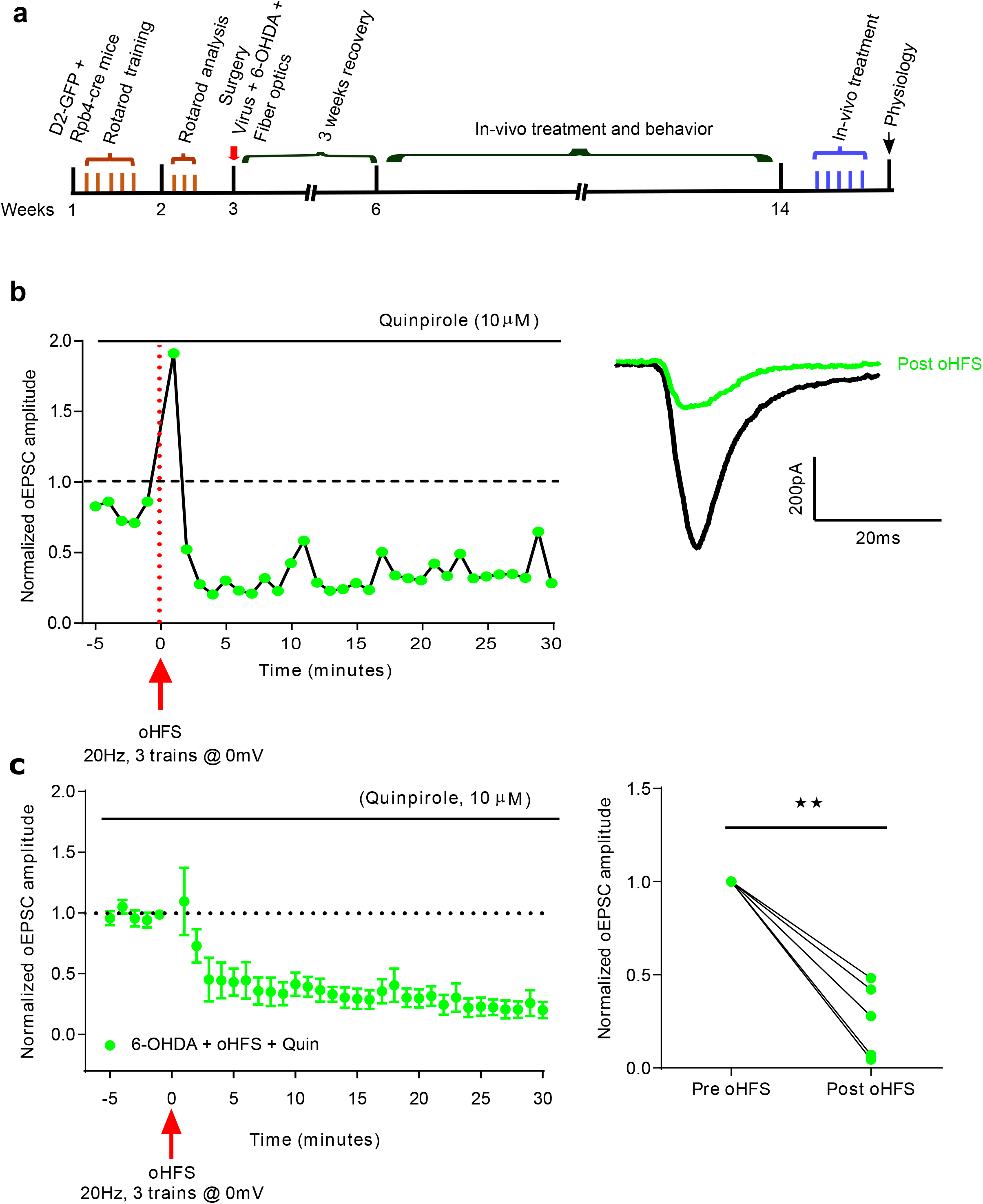
*Ex-vivo* optical stimulation with quinpirole bath application induces LTD in tissue slices from 6-OHDA lesioned animals that received *in-vivo* oHFS/Quin treatment. a. Experimental timeline for brain slice physiology experiments. All the cells recorded within 24 hours of last session of in-vivo oHFS/Quin treatment. b. Representative normalized oEPSCs from a D2-MSN for 5 min recording before oHFS (baseline) and for 30 min after oHFS (left); Raw traces representing baseline (black) and during last 10 min recording (green) of post oHFS (right). c. D2-MSN oEPSCs in DLS were depressed by oHFS (left); all cells showed depression (right) 20 min after oHFS (N=5, from 3 animals **p=0.0028)

